# Rainfall and other meteorological factors as drivers of urban transmission of leptospirosis

**DOI:** 10.1101/658872

**Authors:** Marcelo Cunha, Federico Costa, Guilherme S. Ribeiro, Marília S. Carvalho, Renato B. Reis, Nivison N. Júnior, Lauren Pischel, Edilane L. Gouveia, Andreia C. Santos, Adriano Queiroz, Elsio A. Wunder, Mitermayer G. Reis, Peter J Diggle, Albert I. Ko

**Affiliations:** Escola Nacional de Saúde Pública, Fundação Oswaldo Cruz, Ministério da Saúde, Rio de Janeiro, Brazil; Instituto de Pesquisas Gonçalo Moniz, Fundação Oswaldo Cruz, Ministério da Saúde, Salvador, Brazil; Instituto de Saúde Coletiva, Universidade Federal da Bahia, Salvador, Brazil; Faculty of Health and Medicine, University of Lancaster, Lancaster, UK; Department of Epidemiology of Microbial Diseases, School of Public Health, Yale University, New Heaven, USA; Faculdade de Medicina, Universidade Federal da Bahia, Salvador, Brazil

**Keywords:** Leptospirosis, incidence, seasonal variation, urban, slum, rain, humidity, temperature, time-series, meteorology, climate

## Abstract

**Background:** Leptospirosis is an important public health problem affecting vulnerable urban slum populations in developing country settings. However, the complex interaction of meteorological factors driving the temporal trends of leptospirosis remain incompletely understood.

**Methods and findings:** From 1996 to 2010, we investigated the association between the weekly incidence of leptospirosis and climatic variables in the city of Salvador, Brazil by using a dynamic generalized linear model that accounted for time lags, overall trend and seasonal variation. Our model showed an increase of leptospirosis cases associated with rainfall, lower temperature and higher humidity. There was a lag of one-to-two weeks between weekly values for significant meteorological variables and leptospirosis incidence. Independent of the season, a weekly cumulative rainfall of 20 mm increased the risk of the leptospirosis by 10% compared to a week without rain. Finally, over the 14 year study period the incidence of leptospirosis decreased significantly by four fold (12.8 versus 3.6 per 100,000 people), independently of variations in climate.

**Conclusions:** Strategies to control leptospirosis should focus on avoiding contact with contaminated sources of *Leptospira* as well as on increasing awareness in the population and health professionals within the short time window after both low-level and extreme rainfall events. Increased leptospirosis incidence was restricted to one-to-two weeks after those events suggesting that infectious *Leptospira* survival may be limited to short time intervals.

**Author Summary:** To determine the role of meteorological variables, seasonal variation and temporal trends in the incidence of leptospirosis, we investigated the time series of leptospirosis incidence amongst residents of Salvador, Brazil, from 1996 to 2010. Exploratory and confirmatory statistical methods detected associations between meteorological factors and disease incidence. Results showed the importance of extreme meteorological conditions, particularly rainfall, as short-term predictors of leptospirosis incidence. In addition, we found a long-term decreasing trend of in disease incidence over the observation period.

## INTRODUCTION

Approximately one billion people worldwide reside in slums and this population continues to grow [1]. In those overcrowded environments with inadequate infrastructure, the burden of environmentally-transmitted infections – such as helminths, bacterial diarrheal diseases, leptospirosis among others – is growing and causes more than half a million deaths and 57 million more than half a million deaths and 57 million Disability Adjusted Life-Years annually [2–6]. For these environmentally driven diseases, meteorological factors have a complex interaction with the underlying inadequate sanitation and poor environment influencing the transmission dynamics.

Urban leptospirosis epidemiology is an outstanding example of how the interaction between environment and weather affects human health. More than one million human cases and 58 thousand deaths from leptospirosis are estimated annually around the globe [7]. A large fraction of the global burden of leptospirosis falls on urban slum communities where people live in close proximity to animal and environmental reservoirs of infection [8]. In those settings, leptospirosis transmission is related to the peridomestic household environment [9–12] contaminated with urine from animal reservoirs, most notably rats [13, 14].

Extreme weather events including heavy rainfall and flooding have been consistently associated with an increased incidence of urban leptospirosis [15–17]. In addition to these large sporadic events, seasonal leptospirosis pattern, associated to cyclic rainfall, in endemic areas, have been described in several tropical developing countries [18–25]. Previous studies have suggested that, in addition to rainfall, other climatic factors, such as temperature and humidity may also independently contribute to leptospirosis transmission [18, 20, 26]. A recent study by Matsushita *et al* [27] describes correlations between moderate-to-low rainfall events and clinically suspected cases of leptospirosis, but without considering other climactic factors related to leptospirosis in the urban setting of Manilla. No study to date has disentangled the complex interaction and short time lag of rain, temperature and humidity on the epidemiology of urban leptospirosis. The lack of long-term high quality datasets with appropriate clinical laboratorial case ascertainment, and the low temporal resolution of previous studies [18, 20], have hampered the understanding of the natural history of urban leptospirosis.

Here we establish a relationship between meteorological variables (temperature, rainfall, humidity) and weekly leptospirosis incidence by evaluating data from an active population-based surveillance system for leptospirosis in the Brazilian city of Salvador from 1996 to 2010. We used a dynamic generalized linear model [28] of incidence that took into account the autocorrelation structure of the weekly incidence series to analyze the effects of lag times of meteorological variables, as well as trend and seasonal variation in leptospirosis.

## METHODS

### Study site

Salvador, fourth largest city in the country - is located in north-eastern Brazil and the capital of the state of Bahia. In Brazil notification of leptospirosis cases is mandatory. At the time of this study period, according to the State Secretary of Health’s protocol, patients suspected of leptospirosis are referred to the state infectious diseases hospital (Hospital Couto Maia) for diagnosis and clinical management. This hospital reported 75.4% of the 1,524 reported cases of leptospirosis among residents of Salvador between 2000 and 2010 [29].

### Surveillance

The Oswaldo Cruz Foundation, in collaboration with the Salvador and Bahia Secretaries of Health, has conducted continuous enhanced hospiatal-based surveillance for leptospirosis in Salvador since 1996. Surveillance is primarily based at the state infectious diseases hospital where the study team prospectively evaluated admissions between March 21, 1996 to March 20, 2010 to identify cases who meet the clinical definition for suspected severe leptospirosis, defined as a hospitalized patient with acute undifferentiated fever associated with either bleeding, acute renal failure, jaundice, or acute liver injury with transaminases <1,000 U/L [2][30].

### Laboratory diagnosis of leptospirosis

Study participants had an early acute-phase blood sample collected at enrolment, a late acute-phase blood sample collected 2-7 days later, and a convalescent-phase blood sample collected more than 14 days after the first sample. Blood samples were processed, and sera frozen at −20°C and at −70°C. All sera underwent an IgM enzyme-linked immunosorbent assay (ELISA) (Biomanguinhos, Oswaldo Cruz Foundation, Brazil) and microagglutination test (MAT) using a serum panel comprising a local isolate – *Leptospira interrogans* serovar Copenhageni, strain Fiocruz L1-130 [2] and six reference strains as previously described [31]. *Leptospira* culture was also attempted from acute-phase blood sample [2].

We defined a confirmed case of leptospirosis as one that met at least one of the following criteria: *Leptospira* isolation in blood culture, MAT seroconversion (defined as an increase in MAT titer from 0 to ≥1:200) or MAT four-fold titer rise between paired sera, MAT titer ≥1:800 in a single sample, or a positive result in the IgM-ELISA.

### Meteorological and population data

We obtained climatic data from the meteorological station located in Ondina, Salvador. Data comprised daily measures of rainfall, maximum temperature and relative humidity. The population of Salvador was obtained from a population count in 1996 and from the censuses of 2000 and 2010 [32, 33]. Population counts for the remaining study years were obtained by interpolation from the 1996, 2000 and 2010 data.

### Statistical Analysis

For this dataset, the exact date of admission of leptospirosis cases would give a spurious precision to the time of infection and onset of symptoms, given the imperfect link between infection, onset of symptoms, and hospital admission. Therefore, the number of leptospirosis cases was aggregated per study week, based on the registered date of onset of symptoms. The resulting time series spanned 733 weeks. Various metrics of meteorological data were similarly aggregated into weeks, including weekly accumulated rainfall, weekly average of relative humidity and weekly average of maximum temperature. Three extreme values of rainfall were considered outliers and were truncated at 300mm, to guard against their disproportionately influencing results.

In an exploratory analysis to investigate patterns of overall trend and seasonality, separate working models were fitted to the time series of the transformed incidence, log (incidence + 1), and of the three meteorological variables: weekly accumulated rainfall, weekly average of relative humidity, and weekly average of maximum temperature (see S1 Text). Time-lagged associations between meteorological variables and disease incidence were identified with cross-correlograms of the transformed incidence and residuals of the fitted long-term trend and seasonal effects models for the three meteorological variables. Residual series were used to avoid detection of trivial associations that were a consequence of seasonal variation inherent within all four series.

To investigate the relationship between transformed incidence and explanatory variables, we fitted non-parametric models for the regression of the transformed incidence residual on each of the meteorological residual series using generalized additive models [34].

For the final analysis, we fitted a dynamic generalized linear model [28, 35] in which weekly incidences are assumed to form an independent sequence of Poisson-distributed random variables, conditional on a log-linear multiple regression model that considered meteorological components lagged at 1 and 2 weeks and dynamic (i.e. stochastically varying, trend and seasonal components to capture the correlation structure of the data). The changing size of the population at risk over the study period was treated as an offset in the model (S1 Text).

We applied a Bayesian approach for the estimation of the model parameters using integrated nested Laplace approximations [36] to perform the required calculations. We used the Deviance Information Criterion (DIC) to compare the fits of different models to our data [37]. For each one of these measures, there is no absolute benchmark for a well-fitting model, but in relative terms a smaller value represents a better fit. Analysis was done using *R* (R Development Core Team) [38].

### Ethics Statement

The study protocol received IRB approval from the Hospital Couto Maia, the Oswaldo Cruz Foundation, the Brazilian National Commission for Ethics in Research and Yale University. All adult participants provided written informed consent. Participants aged <18 years and who were able to read signed an informed assent with their parents providing a signed consent. The study team used a standardized data entry form to interview the patients or a family member and to collect data from their medical charts after formal consent. Data included demographics, signs and symptoms at presentation and during hospital stay, clinical management, and disease outcome.

## RESULTS

The surveillance team identified 2,074 clinically suspected cases of leptospirosis. Patients had a mean age of 35 years [Standard deviation (SD): 15] and 1,780 (86%) were men. The mean period between onset of symptoms and hospital admission was 6 days (SD: 3). Clinical characteristics included conjunctival suffusion (637/1,421; 45%), jaundice (1,804/2,074; 87%), renal failure defined by serum creatinine ≥1.5 mg/dL (882/2,074; 42%), and pulmonary haemorrhage with acute lung injury (57/2,074; 3%). Of the 2,074 clinically suspected cases of leptospirosis, 1,498 (72.2%) were laboratory-confirmed. Demographic and clinical presentation of patients were similar between laboratory-confirmed and clinical non-confirmed suspected cases; except for the fatality ratio that was lower among the laboratory-confirmed cases (7%) when compared with clinically suspected non-confirmed cases (25%). Hence, clinically suspected cases were used for further analysis. Statistical analysis using only laboratory confirmed cases resulted in similar findings to those described below (S2 Text).

The time series of weekly incidence shows a pronounced seasonal variation coupled with a decreasing trend (Figure 1). A visual comparison of each of the meteorological time series with the transformed incidence series suggests that the periods of relatively high incidence of leptospirosis, which occur between April and July each year, coincide with periods of high rainfall, high relative humidity and low relative temperature. The cross-correlograms of the incidence residual series with each of the meteorological residual series confirm these associations. They show pronounced peaks (positive for rainfall and humidity, negative for temperature) at one-week and two-week time lags (S1 Fig). Examination of the fitted non-parametric regressions and their associated pointwise 95% confidence limits suggested only a weak departure from linearity between the transformed incidence residual and meteorological residual series (S2 Fig). Together, these results of the exploratory analysis justify our treatment of the one-week and two-week lagged meteorological variables as fixed effects in a log-linear model for incidence.

**Figure 1.**
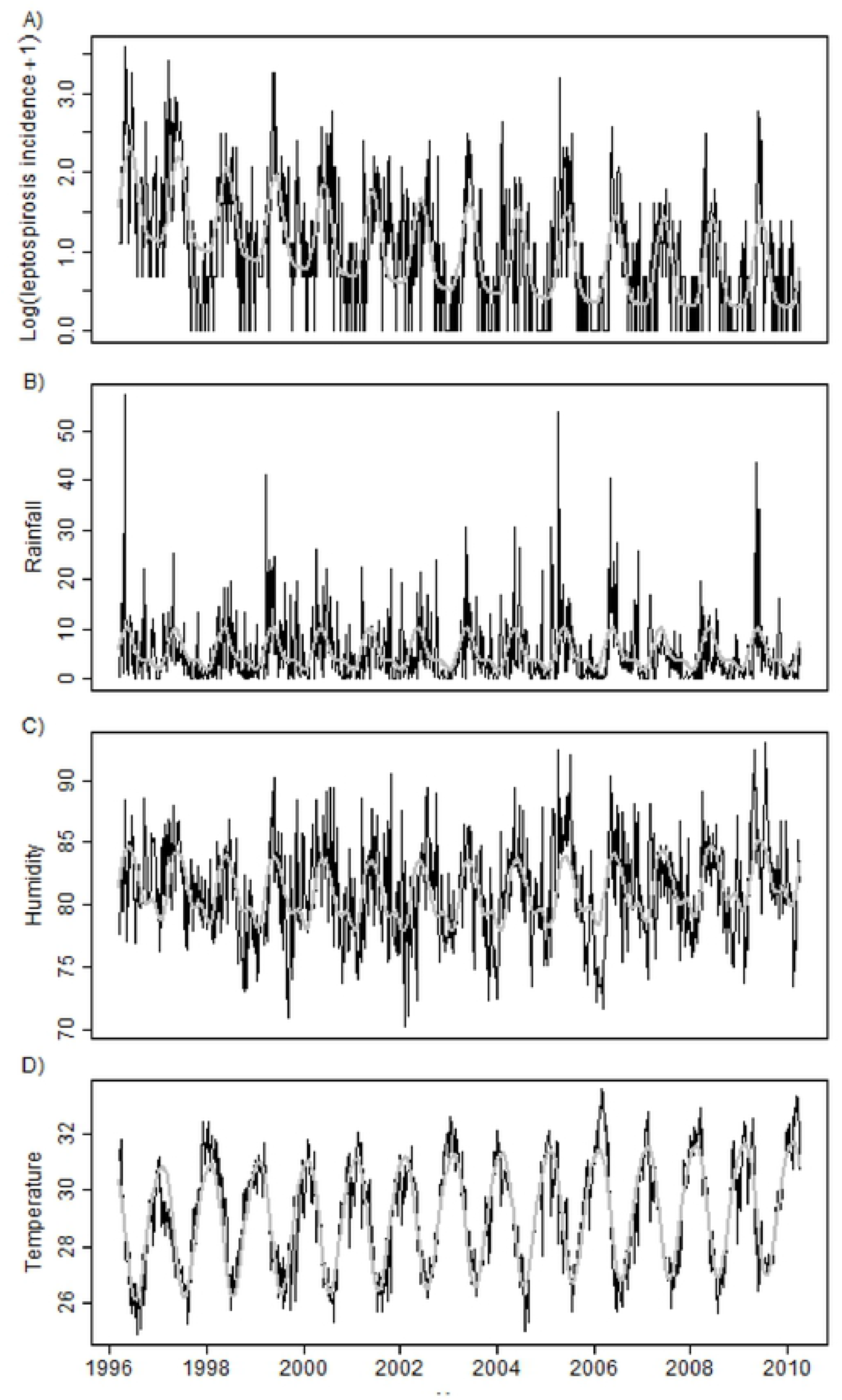
Observed time series (black) and fitted trends (grey) for transformed weekly leptospirosis incidence series (panel A) and meteorological time series: weekly accumulated rainfall in mm (panel B); weekly average of mean daily relative humidity in percent (panel C); and weekly average of mean daily maximum temperature in ° C (panel D), Salvador, Brazil, 1996-2010.

As a first step in the confirmatory analysis, we considered which of the three meteorological variables should be included in the dynamic model. Table 1 compares the DIC values for the eight sub-models that included each possible combination of the three pairs of one-week and two-week lagged meteorological variables. The first four lines show that rainfall was the most important meteorological variable associated with leptospirosis incidence. In addition, we selected the model including all climatic variables that showed the smallest DIC.

**Table 1:**
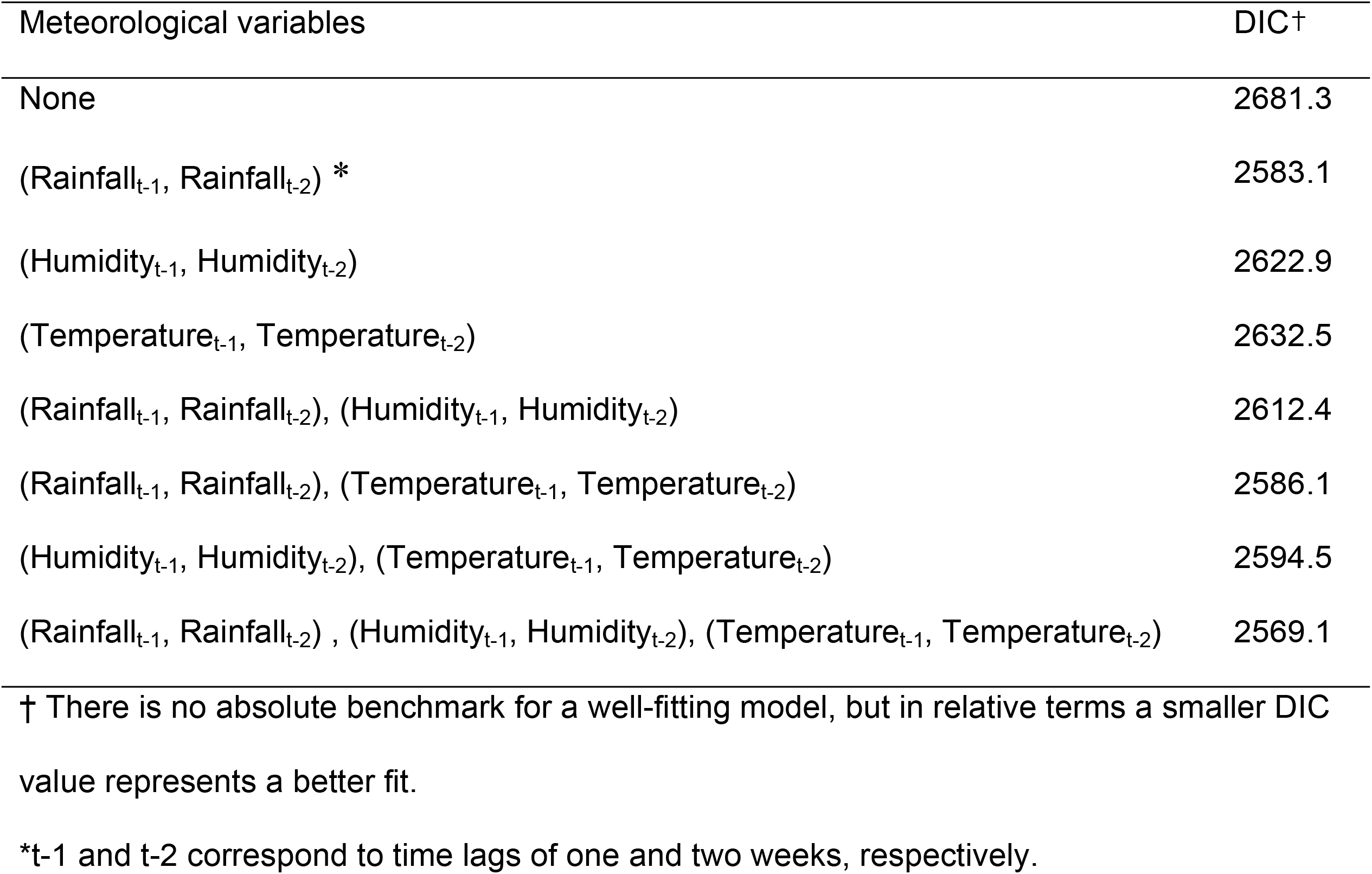
Model fit from eight different combinations of the three pairs of meteorological explanatory variables to predict leptospirosis weekly incidence, Salvador, Brazil, 1996-2010.

| Meteorological variables | DIC† |
| --- | --- |
| None | 2681.3 |
| (Rainfall<sub>t-1</sub>, Rainfall<sub>t-2</sub>) * | 2583.1 |
| (Humidity<sub>t-1</sub>, Humidity<sub>t-2</sub>) | 2622.9 |
| (Temperature<sub>t-1</sub>, Temperature<sub>t-2</sub>) | 2632.5 |
| (Rainfall<sub>t-1</sub>, Rainfall<sub>t-2</sub>), (Humidity<sub>t-1</sub>, Humidity<sub>t-2</sub>) | 2612.4 |
| (Rainfall<sub>t-1</sub>, Rainfall<sub>t-2</sub>), (Temperature<sub>t-1</sub>, Temperature<sub>t-2</sub>) | 2586.1 |
| (Humidity<sub>t-1</sub>, Humidity<sub>t-2</sub>), (Temperature<sub>t-1</sub>, Temperature<sub>t-2</sub>) | 2594.5 |
| (Rainfall<sub>t-1</sub>, Rainfall<sub>t-2</sub>) , (Humidity<sub>t-1</sub>, Humidity<sub>t-2</sub>), (Temperature<sub>t-1</sub>, Temperature<sub>t-2</sub>) | 2569.1 |

† There is no absolute benchmark for a well-fitting model, but in relative terms a smaller DIC value represents a better fit.
\*t-1 and t-2 correspond to time lags of one and two weeks, respectively.

Table 2 shows the estimated change in the relative risk associated with a unit change in each meteorological variable and associated 95% credible intervals. As an example, a weekly cumulative rainfall of 20 mm increased the risk of the leptospirosis by 10% compared to a week without rain (Relative Risk: 1.005; 95% credible interval: 1.004-1.006). All of the interval estimates excluded the neutral value 1, except for one-week-lagged temperature. We preferred to retain this variable because a model that included two-week lagged temperature but excluded one-week lagged temperature would be inconsistent with the known incubation period for the disease (5-14 days) [5].

**Table 2:** Estimated changes in risk of leptospirosis, Salvador, Brazil, 1996-2010. Each row gives the estimate of the relative risk associated with a unit change in the corresponding meteorological variable and the associated 95% Bayesian credible interval.

| Meteorological variable | Relative risk | Credible Interval (95%) |
| --- | --- | --- |
| Rainfall<sub>t-1</sub> | 1.005 | (1.004;1.006) |
| Rainfall<sub>t-2</sub> | 1.003 | (1.002;1.005) |
| Humidity<sub>t-1</sub> | 1.031 | (1.012; 1.050) |
| Humidity<sub>t-2</sub> | 1.041 | (1.022; 1.059) |
| Temperature<sub>t-1</sub> | 0.981 | (0.906;1.062) |
| Temperature<sub>t-2</sub> | 0.881 | (0.816;0.952) |

\*t-1 and t-2 correspond to time lag of one and two weeks, respectively.

The expected weekly incidence according to the fitted model can be decomposed into trend, seasonal and meteorological multiplicative components (Figure 2). Together, these explained about 75.4% of the variation in the observed leptospirosis incidence. The fitted values closely followed the observed temporal distribution of leptospirosis cases (Fig. 2A); 98.1% of the observed values were covered by the 95% credible intervals of the fitted model (not shown). The long-term trend throughout the observation period (Fig. 2B) showed a general decrease in the expected incidence, with a six-fold change between the peak in 1997 and the average of incidence for the period between 2005 and 2010 (1.12 cases per week). For the laboratory confirmed cases we observed an increase after phase 1998 and a decrease phase in incidence since 2002 (S2 Text). These represent temporal changes in expected incidence that were not explained either by seasonal effects or by variation in the residual meteorological series. The seasonal component (Fig. 2C) showed the same characteristic shape in all 14 years, with amplitude corresponding to an approximate doubling of the expected number of cases per week between peak and trough levels, and only slight year-to-year variation in amplitude and phase. The meteorological component (Fig. 2D) demonstrated the importance of extreme meteorological conditions, showing relative long periods close to one, interspersed with sharp peaks in incidence representing up to a five-fold increase.

**Figure 2.**
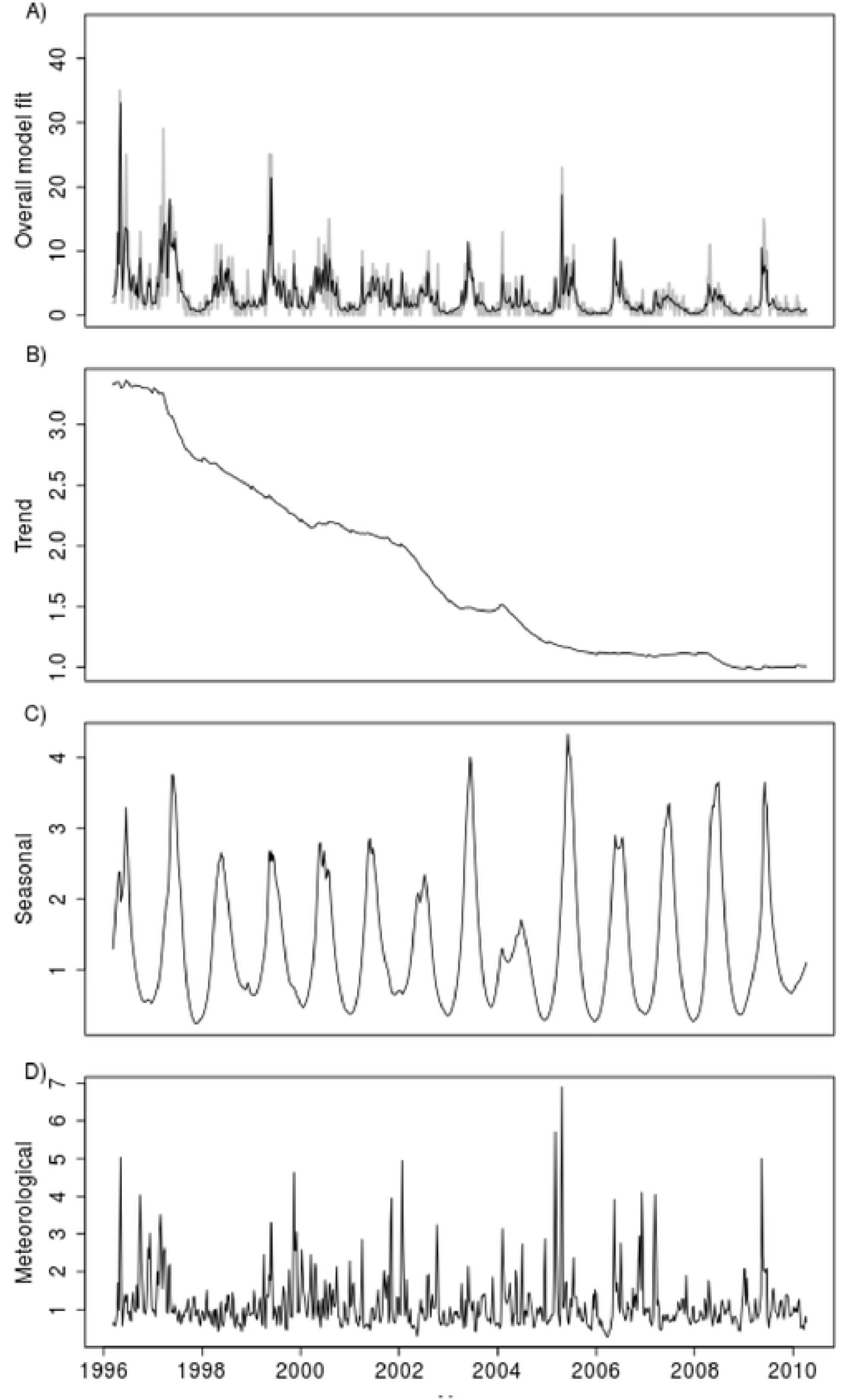
Fitted weekly leptospirosis incidence (black solid line) and observed weekly leptospirosis incidence (grey solid line), and its respective decompositions into trend (panel B), seasonal (panel C) and meteorological (panel D) components, Salvador, Brazil, 1996-2010.

The observed incidence series together with one-week-ahead point forecasts and 95% credible intervals over the last three years (156 weeks) of the observation period are shown in Figure 3. The predictions were well calibrated, as the 95% credible intervals covered 97.4% of the actual values. Note that the credible intervals shown here were for observed rather than expected incidence, i.e. they included an allowance for Poisson variation in each observed count conditional on the underlying risk. Sensitivity analysis showed a relatively good performance for predictions. For example, when the true observed cases were zero, 67% and 97% of predicted values were less than 1 and 2, respectively, whereas when the number of observed cases was 4 or more, about 41% and 88% of the predicted values were greater than 3 and 2 respectively.

**Figure 3.**
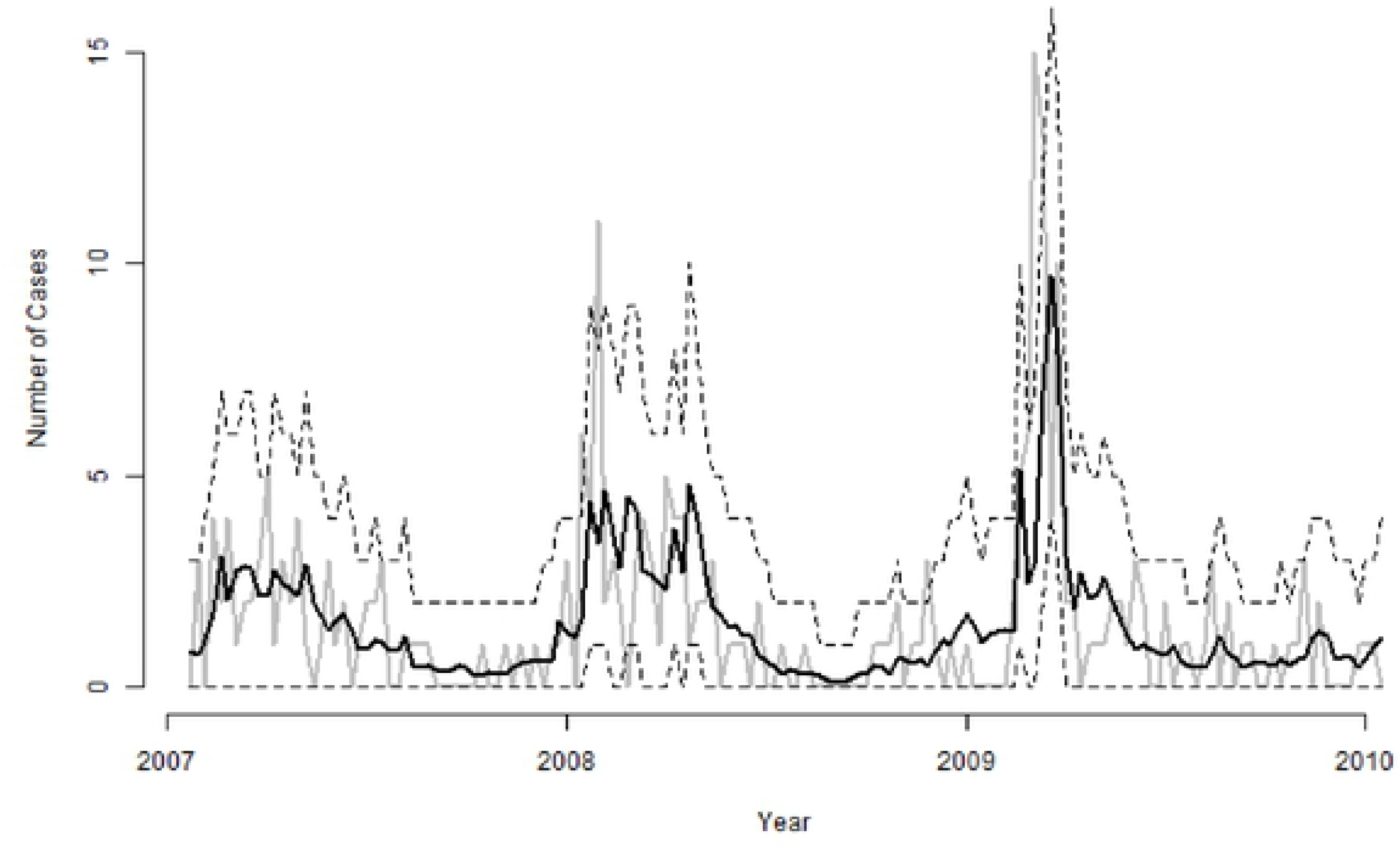
Observed weekly leptospirosis incidence (grey solid line), one-week-ahead forecasts (black solid line) and 95% credible intervals (black dashed lines) for the last three years (2007-2010) of the observations period in Salvador, Brazil.

## DISCUSSION

Leptospirosis is an emerging infectious disease problem in urban slums of developing tropical countries that is driven by two major factors: poverty and climate. This time-series analysis provides novel insights into the meteorological determinants of urban leptospirosis. Using weekly data points, we established and quantified a short-term association between leptospirosis incidence and meteorological variables while taking into account seasonal variation and long-term trend. We demonstrated that a combination of both low-level and extreme rainfall events, as well as temperature and humidity, influence leptospirosis incidence. Increased transmission risk was also restricted to one-to-two weeks after such events, suggesting that the survival of the infective pathogen in the environment may be limited to short time interval. This increased understanding of the natural history of leptospirosis can aid in focusing control measures and support the development of infrastructure that will prevent small scale flooding.

When controlling for seasonal and trend variation, we found an exponential relationship between leptospirosis incidence and meteorological variables. The incidence of leptospirosis cases increased with both low levels of rainfall and extreme rainfall events. This was true even outside the rainy season. For example, when a weekly cumulative rainfall reaches 20 mm (a level frequently observed during the dry season), the risk of the disease increases by 10% compared with a week without rain. While extreme rainfall events affect large areas [39–41], the impact of moderate rainfall on leptospirosis transmission may specifically reflect the precarious state of drainage systems in slum communities, which in turn may increase the risk for exposures to contaminated flood and sewage water. A study from urban Manila in the Philippines also identified that weekly rainfall (but with a different lag time) was associated with increased hospital admission for leptospirosis with the greatest risk after heavy to torrential rainfall. Our data, together with the previous work in Manila, suggest that weekly intensity of rainfall may be a better predictor of leptospirosis incidence than accumulated monthly values, which have been used in previous studies [20, 42, 43].

Our study identified positive short-term associations of rainfall with leptospirosis at lags of 1 and 2 weeks. Previous work in Manila identified that rainfall was associated with higher number of clinically suspected leptospirosis cases at a lag time of 2 weeks [27]. Reasons for the shorter time lags found in our work may be explained by differences in methodology. Our study performed active case finding across the study period for ascertainment of clinical leptospirosis, and our findings were confirmed with the gold standard laboratory testing. The study in Manila included only suspected leptospirosis cases [27] with an expected rate of laboratory confirmation of 54% [44]. Correct estimates of this time lag between rainfall and disease incidence are important for accurate prediction of short-term leptospirosis risk and accurately direct public health interventions. The Bayesian approach used here has the advantage that forecasts of incidence allow for the statistical uncertainty in estimated model parameters.

Our identification of one-to-two-week time lags is consistent with the expected incubation period plus the time until onset of illness, between 5 and 14 days [45]. The 1 to 2 weeks lag between rainfall, temperature, or humidity and leptospirosis identified here, as well as the 2 weeks lag between rainfall and flooding reported in Manila, contrast with results from previous studies that used monthly data and suggested longer temporal association between onset of symptoms of leptospirosis and meteorological data (2 to 10-month lags respectively) [18-20, 46]. Therefore, though leptospires survive for up to one year and remain virulent in culture conditions that mimic the external environment *in vitro* [45, 47, 48], *L. interrogans* survival in spring water decreases to 3 weeks [49]. Clinical infectiveness appears to be in the 1 to 2 weeks after any rainfall event.

In our ananalysis, previous weekly temperatures were negatively associated with leptospirosis incidence while previous weekly humidity was positively associated with disease incidence. The seasonal variations in rainfall and humidity were inversely related to the seasonal temperature variation. A similar relationship was found in Reunion Island [20]. Although humidity and rainfall were moderately correlated (r=0.48) our results suggest that rainfall and humidity have independent effects on disease risk. Both lower temperatures and higher levels of humidity could increase bacterial environmental survival by slowing leptospire desiccation [45] in turn increasing the risk for human leptospirosis [50]. Salvador is a coastal city and part of its relative humidity is dependent, in addition to rainfall, on water evaporation from the Atlantic Ocean. This additional source of humidity could be important for leptospirosis survival in the environment.

Maximum temperature in Salvador varied between 26 and 32°C (Figure 1); changes within this range could affect leptospire survival given that, under laboratory conditions, best growth is obtained between 28-30°C [20, 45]. Our findings differ from results obtained in Reunion Island and Thailand where temperature was positively associated with leptospirosis incidence [18, 20]. However, both of these data sets used mean temperature as opposed to maximum and, in Thailand, monthd with higher temperatures also have higher rainfall. In Salvador, where higher temperatures are registered during the drier months, increased maximum temperature likely increases desiccation, so decreasing leptospire viability in the environment.

There have been limited critical evaluation of statistical approaches on the study meteorological factors and leptospirosis or other water-associated diseases [51]. Previously leptospirosis incidence has been modeled with ARIMA models [18, 20]. Whereas ARIMA require a preliminary transformation of the data to achieve stationary, the dynamic model used in this study can be applied to non-stationary time series without requiring a preliminary transformation of the data. Recently, data from Manila in the Philippines considered distributed lag non-linear models (DLNM) to evaluate the nonlinear temporal dependence between the incidence of leptospirosis and rainfall levels allowing for time delayed effects. Although DLNM has been a useful tool for understanding association of environmental factors and health outcomes [52], it does not account for the time-series data correlation. The dynamic model used here also takes into account for possible nonlinear and lags of time effects between the meteorological variables and the leptospirosis incidence rates. Furthermore, the dynamic model considers the intrinsic temporal data correlation amongst weekly incidences, so that we can be more confident about the validity of our inferences. Additionally, the proposed model makes it possible to evaluate separately the effects for trend, seasonality, and meteorological variables.

We also identified a temporal decline in case incidence that was independent of variations in climate. The modeling approach was useful in evaluating the cumulative influence of interventions designed to reduce disease risk over the study period while controlling for meteorological confounding. There was a significant decline in incidence of severe leptospirosis during the 14-year period with annual incidence four-fold lower in 2009 than in 1996 (3.6 vs. 12.8 per 100,000 population). This decline did not appear to be due to decreased case ascertainment or change in referral patterns. Capture-recapture analyses with national and local databases on hospitalizations, deaths and case reports found that the completeness of the surveillance system was greater than 96%. However, a major intervention was made in the sanitation infrastructure of Salvador during the surveillance period. A $440 million Inter-American Development Bank-supported program (Bahia Azul) constructed 2,000km of closed sewage systems between 1996 and 2004 [53], increasing the city’s population served by safe sewage connections from 26% to 74% [54]. A prospective study demonstrated that the program led to a 22% reduction in diarrheal disease prevalence [53]. Additionally, during 1995-2006 the increase in gross domestic product and social conditional cash transfer programs for the poorest populations in Brazil contributed to reduce poverty and marginally decrease social and economic inequalities [55]. These actions to reduce poverty may have influenced the observed decrease of leptospirosis incidence.

Correlation of seasonal variations in leptospirosis disease with temperature and rainfall does not establish a causal link. Also, we did not include information relating to the spatial distribution of leptospirosis incidence. Our model is related to surveillance of cases and meteorological events within urban Salvador, Bahia and has not been tested for generalizability to other locations or patterns of disease transmission. However, 33% 28% of the Brazilian population live in urban slums [56] with environmental and socioeconomic conditions similar to the lums of Salvador. These figures are comparable to other tropical developing countries [57]. Consequently, we believe that this predictive model could be used, with appropriate local recalibration, in many other urban communities.

This paper highlights important implications for the design of interventions against leptospirosis infection in urban slum communities. We have found that increased risk is associated with short-term, small amounts of rainfall as well as with extreme events. This supports the taking of precautionary measures during any rainy period. Sanitation should be improved in vulnerable communities to prevent even small-scale flooding events. Moreover, we have clearly identified that the period of raised infection risk appears to be during or in the immediate 2 weeks following a rain event. Our models provide a basis for improving early warning systems, and increasing population and medical awareness within this short time window.

## Acknowledgments

We would like to thank the staff of Couto Maia Hospital and the state of Bahia for their assistance in conducting the study. We would like to thank Dr. Jose Hagan for the carefully review of this manuscript. This work was supported by the Coordination for the Improvement of Higher Level Personnel (grant 3373100/2010); the Wellcome Trust (102330/Z/13/Z); and a NSF-NIH grant from the Ecology and Evolution of Infection Diseases (EEID) program (1 R01 TW009504) and NIH grants (5 R01 AI052473, 5 U01 AI088752, 1 R25 TW009338, 1 R01 AI121207, F31 AI114245, FIC D43 TW00919, R01AI121207).

## Supporting Information

**Supplemental Text 1**: Technical appendix for the statistical analysis

**Supplemental Text 2**: Sub-analysis performed for leptospirosis laboratory confirmed cases.

**Supplemental Figure 1**: Cross-correlograms of residuals of transformed leptospirosis incidence with residuals of meteorological variables: weekly accumulated rainfall, weekly average of mean daily relative humidity and weekly average of mean daily maximum temperature, Salvador, Brazil, 1996-2010.

**Supplemental Figure 2:** Generalized additive models for the regression of transformed incidence residuals on weekly accumulated rainfall (left-hand panels), weekly average of mean daily relative humidity (center panels) and weekly average of mean daily maximum temperature (right-hand panels), at one-week (upper row) and two-week (lower row) time-lags, Salvador, Brazil, 1996-2010.

## References

1. UN-HABITAT. Slum Almanac 2015-2016. Tracking Improvement in the Lives of Slum Dwellers. PSUP Team Nairobi. 2016.

2. Ko AI, Galvão Reis M, Ribeiro Dourado CM, Johnson WD, Riley LW. Urban epidemic of severe leptospirosis in Brazil. Salvador Leptospirosis Study Group. Lancet. 1999;354:820–5. doi: 10.1016/S0140-6736(99)80012-9. PubMed PMID: 10485724.

3. Levine MM, Levine OS. Changes in human ecology and behavior in relation to the emergence of diarrheal diseases, including cholera. Proceedings of the National Academy of Sciences of the United States of America. 1994;91(7):2390–4. PubMed PMID: 8146128; PubMed Central PMCID: PMC43377.

4. Bethony J, Brooker S, Albonico M, Geiger SM, Loukas A, Diemert D, et al. Soil-transmitted helminth infections: ascariasis, trichuriasis, and hookworm. Lancet. 2006;367(9521):1521–32. doi: 10.1016/S0140-6736(06)68653-4. PubMed PMID: 16679166.

5. Ko AI, Goarant C, Picardeau M. Leptospira: the dawn of the molecular genetics era for an emerging zoonotic pathogen. Nature Reviews Microbiology. 2009;7(10):736–47. doi: 10.1038/nrmicro2208. PubMed PMID: 19756012; PubMed Central PMCID: PMCPMC3384523.

6. Garchitorena A, Sokolow SH, Roche B, Ngonghala CN, Jocque M, Lund A, et al. Disease ecology, health and the environment: a framework to account for ecological and socio-economic drivers in the control of neglected tropical diseases. Philosophical Transactions of the Royal Society of London B Biological Sciences. 2017;372(1722). doi: 10.1098/rstb.2016.0128. PubMed PMID: 28438917; PubMed Central PMCID: PMCPMC5413876.

7. Torgerson PR, Hagan JE, Costa F, Calcagno J, Kane M, Martinez-Silveira MS, et al. Global Burden of Leptospirosis: Estimated in Terms of Disability Adjusted Life Years. PLoS neglected tropical diseases. 2015;9(10):e0004122. doi: 10.1371/journal.pntd.0004122. PubMed PMID: 26431366; PubMed Central PMCID: PMC4591975.

8. Costa F, Carvalho-Pereira T, Begon M, Riley L, Childs J. Zoonotic and Vector-Borne Diseases in Urban Slums: Opportunities for Intervention. Trends Parasitol. 2017;33(9):660–2. doi: 10.1016/j.pt.2017.05.010. PubMed PMID: 28625886.

9. Sarkar U, Nascimento SF, Barbosa R, Martins R, Nuevo H, Kalafanos I, et al. Population-based case-control investigation of risk factors for leptospirosis during an urban epidemic. Am J Trop Med Hyg. 2002;66:605–10.

10. Reis RB, Ribeiro GS, Felzemburgh RDM, Santana FS, Mohr S, Melendez AXTO, et al. Impact of environment and social gradient on Leptospira infection in urban slums. PLoS neglected tropical diseases. 2008;2:e228. doi: 10.1371/journal.pntd.0000228. PubMed PMID: 18431445.

11. Felzemburgh R, Ribeiro GS, Costa F, Reis RB, Hagan JE, Melendez AX, et al. Prospective study of leptospirosis transmission in an urban slum community: Role of poor environment in repeated exposures to the Leptospira agent. PLoS neglected tropical diseases. 2014;In press.

12. Hagan JE, Moraga P, Costa F, Capian N, Ribeiro GS, Wunder EA, Jr., et al. Spatiotemporal Determinants of Urban Leptospirosis Transmission: Four-Year Prospective Cohort Study of Slum Residents in Brazil. PLoS neglected tropical diseases. 2016;10(1):e0004275. doi: 10.1371/journal.pntd.0004275. PubMed PMID: 26771379; PubMed Central PMCID: PMC4714915.

13. Costa F, Ribeiro GS, Felzemburgh RD, Santos N, Reis RB, Santos AC, et al. Influence of household rat infestation on leptospira transmission in the urban slum environment. PLoS neglected tropical diseases. 2014;8(12):e3338. doi: 10.1371/journal.pntd.0003338. PubMed PMID: 25474580; PubMed Central PMCID: PMC4256176.

14. Costa F, Wunder EA, Jr., De Oliveira D, Bisht V, Rodrigues G, Reis MG, et al. Patterns in Leptospira Shedding in Norway Rats (Rattus norvegicus) from Brazilian Slum Communities at High Risk of Disease Transmission. PLoS neglected tropical diseases. 2015;9(6):e0003819. doi: 10.1371/journal.pntd.0003819. PubMed PMID: 26047009; PubMed Central PMCID: PMCPMC4457861.

15. Gaynor K, Katz AR, Park SY, Nakata M, Clark TA, Effler PV. Leptospirosis on Oahu: An outbreak associated with flooding of a University Campus. American Journal of Tropical Medicine and Hygiene. 2007;76(5):882–5. PubMed PMID: WOS:000246326300019.

16. Pappachan MJ, Sheela M, Aravindan KP. Relation of rainfall pattern and epidemic leptospirosis in the Indian State of Kerala (vol 58, pg 1054, 2004). J Epidemiol Commun H. 2005;59(3):251–. PubMed PMID: WOS:000228010200018.

17. Maskey M, Shastri JS, Saraswathi K, Surpam R, Vaidya N. Leptospirosis in Mumbai: post-deluge outbreak 2005. Indian journal of medical microbiology. 2006;24:337–8. PubMed PMID: 17185871.

18. Chadsuthi S, Modchang C, Lenbury Y, Iamsirithaworn S, Triampo W. Modeling seasonal leptospirosis transmission and its association with rainfall and temperature in Thailand using time-series and ARIMAX analyses. Asian Pac J Trop Med. 2012;5(7):539–46. doi: Doi 10.1016/S1995-7645(12)60095-9. PubMed PMID: WOS:000305107700008.

19. Weinberger D, Baroux N, Grangeon JP, Ko AI, Goarant C. El Nino Southern Oscillation and Leptospirosis Outbreaks in New Caledonia. PLoS neglected tropical diseases. 2014;8(4). doi: DOI 10.1371/journal.pntd.0002798. PubMed PMID: WOS:000335342400033.

20. Desvars A, Jego S, Chiroleu F, Bourhy P, Cardinale E, Michault A. Seasonality of Human Leptospirosis in Reunion Island (Indian Ocean) and Its Association with Meteorological Data. Plos One. 2011;6(5). doi: ARTN e20377 DOI 10.1371/journal.pone.0020377. PubMed PMID: WOS:000291097600059.

21. Hirschauer C, Daudens E, Coudert C, Frogier E, Melix G, Fleure M, et al. Epidemiology of leptospirosis in French Polynesia from 2006 to 2008 [in French]. Bulletin Epidémiologique Hebdomadaire 2009;48-49-50:508–11.

22. Lhomme V, Grolierbois L, Jouannelle J, Elisabeth L. Leptospirose en Martinique de 1987 à 1992: bilan d’une étude épidémiologique, clinique et biologique. Med Mal Infect. 1996;26:94–8. doi: 10.1016/S0399-077X(96)80161-2.

23. Herrmann-Storck C, Brioudes A, Quirin R, Deloumeaux J, Lamaury I, Nicolas M, et al. Retrospective review of leptospirosis in Guadeloupe, French West Indies 1994-2001. West Indian Med J. 2005;54(1):42–6.

24. Tassinari WdS, Pellegrini DDCP, Sá CBP, Reis RB, Ko AI, Carvalho MS. Detection and modelling of case clusters for urban leptospirosis. Tropical medicine & international health. 2008;13:503–12. doi: 10.1111/j.1365-3156.2008.02028.x. PubMed PMID: 18312472.

25. Mohan ARM, Cumberbatch A, Adesiyun AA, Chadee DD. Epidemiology of human leptospirosis in Trinidad and Tobago, 1996-2007: a retrospective study. Acta tropica. 2009;112:260–5. doi: 10.1016/j.actatropica.2009.08.007. PubMed PMID: 19679092.

26. Sumi A, Telan EF, Chagan-Yasutan H, Piolo MB, Hattori T, Kobayashi N. Effect of temperature, relative humidity and rainfall on dengue fever and leptospirosis infections in Manila, the Philippines. Epidemiol Infect. 2017;145(1):78–86. doi: 10.1017/S095026881600203X. PubMed PMID: 27608858.

27. Matsushita N, Ng CFS, Kim Y, Suzuki M, Saito N, Ariyoshi K, et al. The non-linear and lagged short-term relationship between rainfall and leptospirosis and the intermediate role of floods in the Philippines. PLoS neglected tropical diseases. 2018;12(4):e0006331. doi: 10.1371/journal.pntd.0006331. PubMed PMID: 29659576; PubMed Central PMCID: PMC5919665.

28. West M, Harrison J. Bayesian Forecasting and Dynamic Models. 2nd ed: Springer; 1997.

29. Salvador. Secretaria Municiapal de Saúde. http://www.tabnet.saude.salvador.ba.gov.br/. 2010.

30. Nabity SA, Ribeiro GS, Aquino CL, Takahashi D, Damião AO, Gonçalves AHO, et al. Accuracy of a dual path platform (DPP) assay for the rapid point-of-care diagnosis of human leptospirosis. PLoS neglected tropical diseases. 2012;6:e1878. doi: 10.1371/journal.pntd.0001878. PubMed PMID: 23133686.

31. Gouveia EL, Metcalfe J, de Carvalho ALF, Aires TSF, Villasboas-Bisneto JC, Queirroz A, et al. Leptospirosis-associated severe pulmonary hemorrhagic syndrome, Salvador, Brazil. Emerg Infect Dis. 2008;14:505.

32. Estadística IBdGe. Dados do Censo 2010 publicados no Diário Oficial da União do dia 04/11/2010. Censo 2010. 2010.

33. IBGE. Resultados do Universo. Rio de Janeiro: Instituto Brasileiro de Geografia e Estatística. Instituto Brasileiro de Geografia e Estatística. Censo Demográfico 2000. 2002.

34. Hastie T, Tibshirani R, Friedman J. The Elements of Statistical Learning: Data Mining, Inference, and Prediction. Second ed. New York: Springer; 2009.

35. West M, Harrison P, Migon H. Dynamic generalized linear models and Bayesian forecastin. Journal of the American Statistical. 1985;80(389):73–83.

36. Rue H, Martino S, Chopin N. Approximate Bayesian inference for latent Gaussian models by using integrated nested Laplace approximations. J R Stat Soc B. 2009;71:319–92. doi: DOI 10.1111/j.1467-9868.2008.00700.x. PubMed PMID: WOS:000264374200002.

37. Spiegelhalter DJ, Best NG, Carlin BP, van der Linde A. Bayesian measures of model complexity and fit. Journal of the Royal Statistical Society: Series B (Statistical Methodology). 2002;64:583–639. doi: 10.1111/1467-9868.00353. PubMed PMID: 6175412381464386404.

38. RStudio Team. RStudio: Integrated Development for R. RStudio I, Boston, MA URL http://www.rstudio.com/.2015.

39. Agampodi SB, Peacock SJ, Thevanesam V, Nugegoda DB, Smythe L, Thaipadungpanit J, et al. Leptospirosis outbreak in Sri Lanka in 2008: Lessons for assessing the global burden of disease. Am J Trop Med Hyg. 2011;85:471–8. doi: 10.4269/ajtmh.2011.11-0276.

40. Thaipadungpanit J, Wuthiekanun V, Chierakul W, Smythe LD, Petkanchanapong W, Limpaiboon R, et al. A dominant clone of Leptospira interrogans associated with an outbreak of human leptospirosis in Thailand. PLoS neglected tropical diseases. 2007;1:e56.

41. Ledien J, Sorn S, Hem S, Huy R, Buchy P, Tarantola A, et al. Assessing the performance of remotely-sensed flooding indicators and their potential contribution to early warning for leptospirosis in Cambodia. Plos One. 2017;12(7):e0181044. doi: 10.1371/journal.pone.0181044. PubMed PMID: 28704461; PubMed Central PMCID: PMC5509259.

42. Kupek E, de Sousa Santos Faversani MC, de Souza Philippi JM. The relationship between rainfall and human leptospirosis in Florianópolis, Brazil, 1991-1996. Brazilian journal of infectious diseases. 2000;4:131–4.

43. Barcellos C, Sabroza PC. The place behind the case: leptospirosis risks and associated environmental conditions in a flood-related outbreak in Rio de Janeiro. Cad Saude Publica. 2001;17 Suppl:59–67. doi: 10.1590/S0102-311X2001000700014. PubMed PMID: 11426266.

44. Kitashoji E, Koizumi N, Lacuesta TL, Usuda D, Ribo MR, Tria ES, et al. Diagnostic Accuracy of Recombinant Immunoglobulin-like Protein A-Based IgM ELISA for the Early Diagnosis of Leptospirosis in the Philippines. PLoS neglected tropical diseases. 2015;9(6):e0003879. doi: 10.1371/journal.pntd.0003879. PubMed PMID: 26110604; PubMed Central PMCID: PMC4482399.

45. Faine S. Leptospira & Leptospirosis. 1999:368. Epub 1.

46. Joshi YP, Kim EH, Cheong HK. The influence of climatic factors on the development of hemorrhagic fever with renal syndrome and leptospirosis during the peak season in Korea: an ecologic study. BMC Infect Dis. 2017;17(1):406. doi: 10.1186/s12879-017-2506-6. PubMed PMID: 28592316; PubMed Central PMCID: PMC5463320.

47. Trueba G, Zapata S, Madrid K, Cullen P, Haake D. Cell aggregation: a mechanism of pathogenic Leptospira to survive in fresh water. Int Microbiol. 2004;7(1):35–40. PubMed PMID: WOS:000221604600006.

48. Stoddard R, Bui D, Haberling D, Wuthiekanun V, Thaipadungpanit J, Hoffmaster AR. Viability of Leptospira Isolates from a Human Outbreak in Thailand in Various Water Types, pH, and Temperature Conditions. Am J Trop Med Hyg. 13:0748.

49. Casanovas-Massana A, Pedra GG, Wunder EA, Jr., Diggle PJ, Begon M, Ko AI. Quantification of Leptospira interrogans Survival in Soil and Water Microcosms. Appl Environ Microbiol. 2018;84(13). doi: 10.1128/AEM.00507-18. PubMed PMID: 29703737; PubMed Central PMCID: PMC6007094.

50. Levett PN. Leptospirosis. Clin Microbiol Rev. 2001;14(2):296–326. doi: 10.1128/CMR.14.2.296-326.2001. PubMed PMID: 11292640; PubMed Central PMCID: PMC88975.

51. Lo Iacono G, Armstrong B, Fleming LE, Elson R, Kovats S, Vardoulakis S, et al. Challenges in developing methods for quantifying the effects of weather and climate on water-associated diseases: A systematic review. PLoS neglected tropical diseases. 2017;11(6):e0005659. doi: 10.1371/journal.pntd.0005659. PubMed PMID: 28604791; PubMed Central PMCID: PMC5481148.

52. Gasparrini A, Armstrong B, Kenward MG. Distributed lag non-linear models. Stat Med. 2010;29(21):2224–34. doi: 10.1002/sim.3940. PubMed PMID: 20812303; PubMed Central PMCID: PMC2998707.

53. Barreto ML, Genser B, Strina A, Teixeira MG, Assis AMO, Rego RF, et al. Effect of city-wide sanitation programme on reduction in rate of childhood diarrhoea in northeast Brazil: assessment by two cohort studies. Lancet. 2007;370(9599):1622–8. doi: Doi 10.1016/S0140-6736(07)61638-9. PubMed PMID: WOS:000250883400021.

54. AESBE. Associação das Empresas de Saneamento Básico Estaduais - Aesbe. Embasa consegue reduzir custos e assina contrato de PPP. Associação das Empresas de Saneamento Básico Estaduais. http://www.aesbe.org.br/aesbe/pages/noticia/exibirLeitura.do?id=153[WebSite]. 2007.

55. Hoffman R, Kageyama A. Poverty in Brazil: Multidimensional Perspective Economia e Sociedade. 2006;15(1):79–112.

56. IBGE. Subnormal Agglomerates - First Results. In: Estatística IBdGe, editor. Rio de Janeiro 2010. p. 1–259.

57. UN-HABITAT. State of the World’s Cities 2010/2011: Bridging The Urban Divide. GB. Earthscan. 2010.

